# Rhythms and Background (RnB): The Spectroscopy of Sleep Recordings

**DOI:** 10.1101/2024.10.28.620621

**Authors:** J. Dubé, M. Foti, S. Jaffard, V. Latreille, B. Frauscher, J. Carrier, J.M. Lina

## Abstract

Non-rapid eye movement (NREM) sleep is characterized by the interaction of multiple coupled oscillations essential for various functions such as memory consolidation, alongside a pervasive and dynamic arrhythmic 1/f scale-free background that may also contribute to these functions. Although recent spectral parametrization methods such as FOOOF (Fitting-One-and-Over-f) allowed to dissociate rhythmic and arrhythmic components in the spectral domain, they fail to resolve these processes in the time domain, where instantaneous measures of frequency, amplitude, and phase-amplitude coupling are still confounded by arrhythmic activity. This limitation represents a significant pitfall for studies of NREM sleep, which often rely on phase-based analyses of specific oscillations. To address this limitation, we introduce ‘Rhythms & Background’ (RnB), a novel wavelet-based methodology designed to dynamically denoise time-series data by correcting for arrhythmic interference. This enables the extraction of a purely rhythmic time-series suitable for enhanced time-domain analyses of sleep rhythms. We first validate RnB through simulations, demonstrating its robust performance in accurately estimating spectral profiles of individual and multiple oscillations across a range of arrhythmic conditions. We then apply RnB to publicly available intracranial EEG sleep recordings, showing that it provides an improved spectral and time-domain representation of hallmark NREM rhythms. Finally, we demonstrate that RnB significantly enhances the assessment of phase-amplitude coupling between cardinal NREM oscillations, outperforming traditional methods that conflate rhythmic and arrhythmic components. This methodological advance offers a substantial improvement in the analysis of sleep oscillations, providing greater precision in the study of rhythmic activity critical to NREM sleep functions.

**SIGNIFICANCE STATEMENT:** The Rhythms and Background (RnB) algorithm introduces a novel approach to signal processing in electrophysiology by isolating rhythmic activity from the arrhythmic background at the time-series level. Unlike existing spectral decomposition methods, RnB enables more precise analysis of brain rhythms in both the time and spectral domains, providing clearer insights into cerebral oscillatory processes. This breakthrough has direct applications in studying brain connectivity and oscillation dynamics during sleep. Additionally, its application in clinical populations where pathological changes in arrhythmic activity are common, such as neurodevelopmental and neurodegenerative disorders, will help to better understand abnormal oscillatory processes. By improving the accuracy of rhythmic signal analysis, RnB opens new avenues for understanding brain function and dysfunction in research and clinical settings.

## Introduction

Brain rhythms from bioelectric activity reflect synchronization of neuronal assemblies and are often inventoried using the Fourier analysis of electrophysiological brain recordings (Buzsáki, 2004). A noteworthy case is non-rapid eye movement (NREM) sleep, which is characterized by cardinal rhythms critically involved in overnight information processing such as sleep slow waves (SSW, 0.5-4 Hz), theta bursts (6-10 Hz), sleep spindles (8-16 Hz), and sharp-wave ripples (100-200 Hz). These rhythms are organized in complex wave sequences, with sporadic phase-amplitude coupling across frequencies, which occur locally and between remote brain regions (Klinzing et al., 2016; Helfrich et al., 2019). For instance, sleep spindles usually occur after SSWs down states, within the transition period towards their upstate. The loss of specificity of this phase coupling in aging may contribute to impaired memory consolidation (Helfrich et al., 2018).

New hypotheses suggest that arrhythmic brain activity plays complementary roles to brain rhythms in NREM sleep functions (Helfrich et al., 2021). Arrhythmic activity recruits a broad range of frequencies and is usually expressed as 1/*f^β^* decay in the power spectral density (He, 2014). This “scale-free” arrhythmic component is usually estimated using the spectral slope (*β*) expressed in the log-log power spectrum. The steepness of *β* has been shown to closely follow shift in the excitation-inhibition balance during sleep (Lendner et al., 2020; Helfrich et al., 2021; Niethard et al., 2021; Kozhemiako et al., 2022), and to show age-dependent changes which can predict ulterior cognitive impairment (Finley et al., 2024; Hernandez et al., 2024). Furthermore, arrhythmic differences in *β* alter the spectral peaks’ characteristics in the Fourier domain which are typically attributed to brain rhythms (i.e. width, height, central frequency). Spectral parametrization methods, such as *Fitting-Oscillations-One-and-Over-F (FOOOF),* have been recently introduced to robustly assess rhythmic (peaks) and arrhythmic (*β*) components in the power spectrum (Wen and Liu, 2016; Donoghue et al., 2020). Such methods are increasingly used to report rhythmic and arrhythmic changes in neuroscience (Donoghue et al., 2022; Ostlund et al., 2022).

Arrhythmicity is also evident at the signal level, manifesting as a ubiquitous ‘desynchronized’ background activity that nonlinearly interferes with transient oscillatory events (Helfrich et al., 2021). This interplay introduces noise in time-domain analyses, reducing the reliability of phase and amplitude estimates typically associated to brain rhythms (Jurkiewicz et al., 2021; Donoghue et al., 2022). Such metrics are crucial for investigating oscillation-dependent coupling and connectivity in neurodegenerative disorders such as Alzheimer’s and Parkinson’s disease (Nimmrich et al., 2015). Therefore, correcting the phase and amplitude instantaneous estimates for differences in arrhythmicity — which is now increasingly documented across various brain dysfunctions (Pani et al., 2022) — is imperative. However, no existing methods currently address this challenge.

The present work addresses this critical gap by introducing a novel wavelet-based algorithm, **Rhythms & Background (*RnB*)**. This algorithm accounts for the arrhythmic background in NREM sleep’s electrophysiological time series by denoising rhythmic activity at the level of wavelet coefficients in the ‘time scale’ domain. By synthesizing corrected wavelet coefficients, RnB extracts a ’rhythmic time series’ in which arrhythmic activity is attenuated. *RnB* thus allows an accurate representation of brain rhythms, effectively performing a ‘spectroscopy’ of NREM sleep recordings.

The method section begins by presenting the mathematical framework that allows the transition from the spectral to the wavelet paradigm. This section also outlines the key methodological tools—such as a slow-wave detector, time-frequency analyses, and amplitude-phase coupling—employed in subsequent analyses of rhythmic time series derived from NREM sleep recordings. Next, we validate the algorithm’s ability to estimate background and rhythmic components using a simulation paradigm. Following this, we apply the RnB algorithm to intracranial NREM sleep recordings to conduct a detailed spectroscopy and inventory of NREM sleep rhythms. Finally, we demonstrate that the RnB-derived rhythmic time series enable the detection of rhythmic sleep slow waves during NREM sleep, which exhibit robust coupling to other NREM rhythms.

## Materials and Methods

### From the spectral to the wavelet paradigm of rhythms and background

Spectral approaches characterize rhythms and background from the spectral power in the Fourier domain. Without loss of generality, we considered the following model of the power spectral density Γ(*f*)

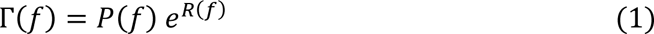

where the factor *P*(*f*) = *c*/*f^β^* accounts for arrhythmicity (as a constant *c* and a slope *β* also called scaling exponent) and *R*(*f*) represents rhythms. Expanding the exponential, this multiplicative model can accommodate the usual additive model,

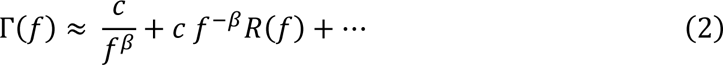

where the first term accounts for the arrhythmic background, and the second term is a resonance, if *R*(*f*) is sufficiently decaying at low frequency to compensate the infrared divergence. This expansion can explain the possible modulation of narrow band power peaks. To extract the arrhythmic factor, we follow the IRASA approach (Wen and Liu, 2016), considering a random sequence of power spectrums obtained from a series of rescaling transforms *f* → *af*,

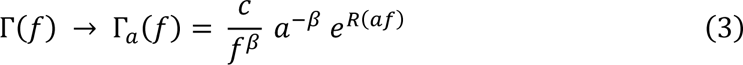

where 0 < *a* < 2 will be chosen in pair, *a* and 1/*a*. Then, from *n* randomly selected values for *a*, the geometric average

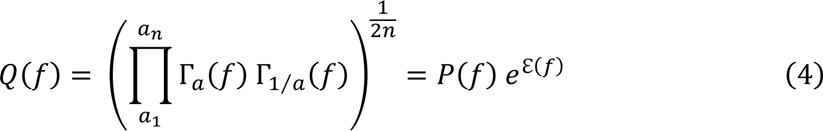

exhibits the arrhythmic factor *P*(*f*) = *c/f^β^* since a large number of random rescaling will not preserve narrowband rhythmic processes present in the spectrum as previously demonstrated, and 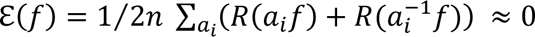

Thus, *Q*(*f*)^−1^ ≈ *P*(*f*)^−1^ is a spectral filter able to whiten the arrhythmic component of the individual power spectrums on an epoch-per-epoch basis. The rhythmic factor *e*^*R*\*(*f*)^ is estimated from the average of this residual spectra across all epochs ’*i*’,

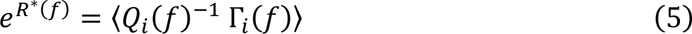

which reliably estimates the spectral signature of simulated oscillations (see Fig.1, E1 and E2). It is worth noting that the spectral slope can be estimated from a log-log regression of (3) applied for each epoch, leading to a spectrum of the scaling exponent as shown in Fig.1(D1 and D2).

**Fig.1.**
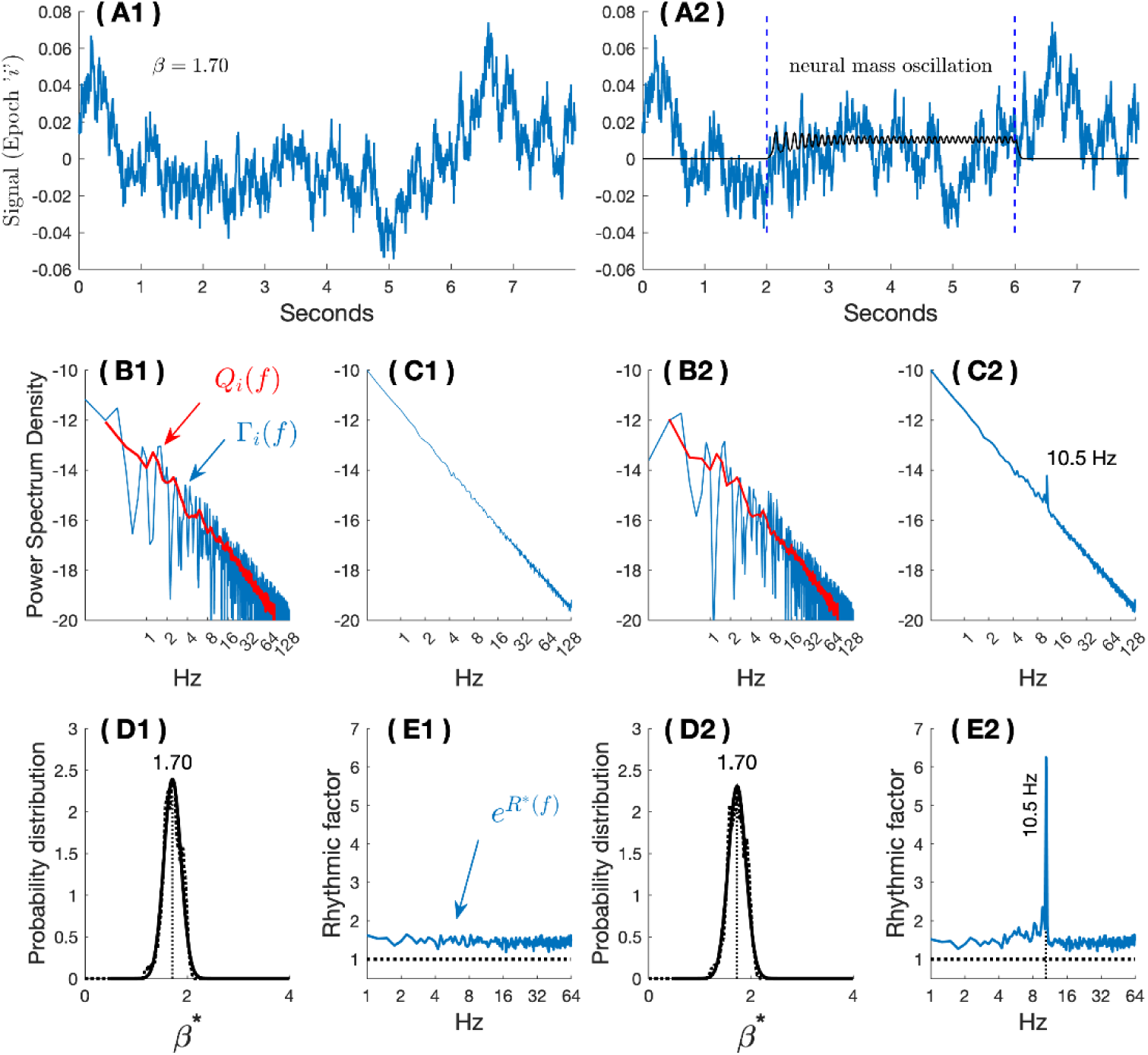
Spectral estimation of arrhythmic background and rhythms. (A1, A2). Examples of two simulated time series (see simulations section for methodological details). (A1) show pure background arrhythmic activity (*β* =1.70; arrhythmic background). (A2) shows a mixture of the same arrhythmic activity as in A1, with the addition of a four second alpha oscillation (10.5 Hz). (B1, B2). Fourier power spectra of (A1) and (A2), respectively. The estimated arrhythmic spectrum *Q_i_*(*f*), as described above, is shown in red. (C1, C2). Averaged standard power spectra over 250 simulations for each type of simulation as shown in (B1) and (B2). Even though the added oscillation in the signal (A2) is unnoticeable in the individual Fourier spectrum (B2), its signature is apparent in the averaged standard spectrum (C2). (D1, D2). Probability density of estimated scale-free exponents obtained from *Q_i_*(*f*) in the two sets of 250 simulations, without (D1) and with (D2) the alpha 10.5 Hz oscillation. (E1, E2). the residual ‘rhythmic factor’ exp(*R*^*^(*f*)) controlled for the presence of arrhythmic background for the two sets of simulations, respectively without (E1) and with (E2) the alpha 10.5 Hz oscillation.

As for all current spectral approaches, this filtering approach cannot restore the underlying rhythmic signal. To disentangle rhythms from arrhythmicity in the time domain, a signal processing approach mixing time *and* frequency is mandatory.

This is precisely the wavelet paradigm developed in *RnB*.

### Wavelet and arrhythmicity

Although the scale-free arrhythmic process is usually addressed in the Fourier power spectrum, the ‘time – scale decomposition’ is an appropriate representation of such a process (Rocca et al., 2018; Abry et al., 2019). This representation amounts to expand the signal in terms of ‘transient little wave’ called wavelets,

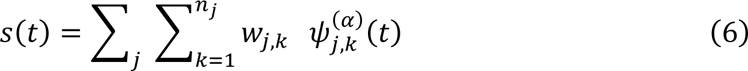

In this expansion, each wavelet has a temporal scale ≈ 2^*j*^ and is timely located at *t* = *k* 2^*j*^ where *k* and *j* are integer,

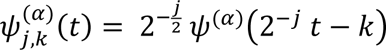

The smoothness of the ‘mother wavelet’ *ψ*^(*α*)^(*t*) is controlled by the regularity parameter *α*. This ‘little wave’ oscillates with a frequency *f*_0_ that depends on the parameter *α*. Therefore, the main frequency of the wavelets at scale *j* in expansion (6) is ≈ 2^−*j*^ *f*_0_.

The multiresolution ‘time - scale’ representation of a signal in such a basis is a collection of wavelet coefficients at various discrete time and scales as is illustrated in Fig.2C.

**Fig.2.**
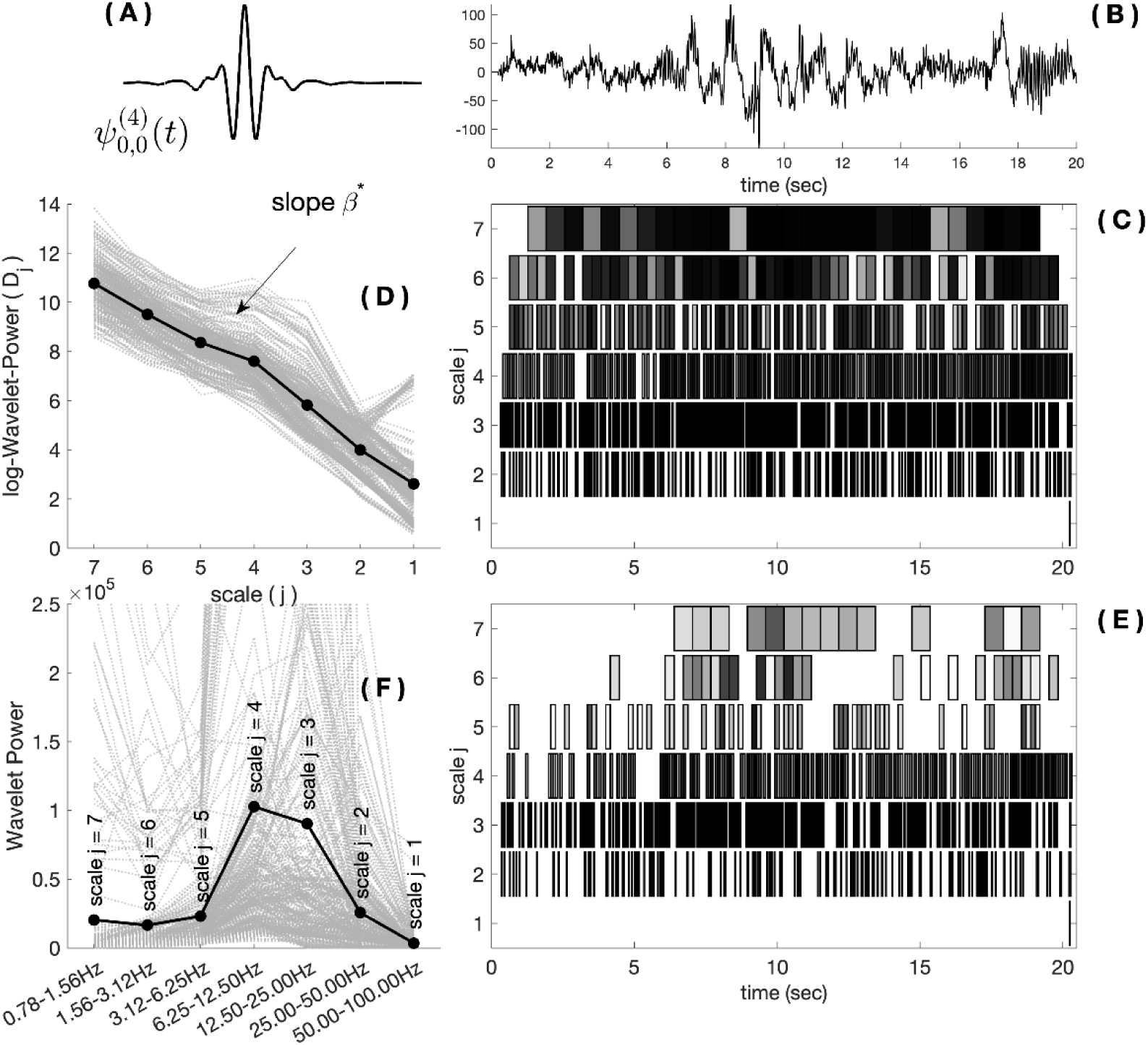
The analysis of an original signal and synthesis of a rhythmic signal using fractional spline wavelets. (A) An example of fractional spline wavelet that could be used for analysis of an NREM sleep signal as illustrated in (B). (C) ‘time-scale’ decomposition of *s*(*t*) displayed in (B) using wavelet basis generated with (A). Each boxes represents a wavelet coefficient 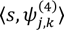 at a particular scale (*j*) and discrete time (*k* 2^*j*^). Real time is indicated on the x-axis. To help the reading of the time-scale representations, visible boxes account for 99% of the overall wavelet power of the signal. Gray intensity (globally normalized) indicates the amplitude of the largest wavelet coefficient. (D) Logarithmic wavelet power of (B) with respect to the dyadic scale parameter. The linear regression will provide the scaling exponent as given in (6). (E) ‘time-scale’ decomposition of the rhythmic signal resulting from processing (B). Notice the lacunar representation as compared to the analysis of the original signal in (C). The rhythmic decomposition exhibits coefficients at large scale (low frequencies) that seem to account for oscillatory pattern localized in time (confirmed with an inspection of (B)). (F) Wavelet power of multiple rhythmic signals (dashed grey lines) and the mean (dark bold line).

The previous log-log regression allowing to estimate the scaling exponents in the spectral domain, log Γ(*f*) ≈ −*β* log *f*, can be reformulated in the wavelet domain as follows

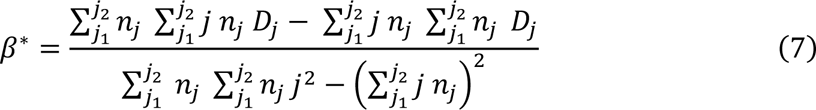

with the second-order statistics of the wavelet coefficients,

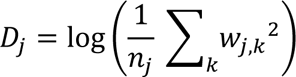

The regression, illustrated on Fig. 2D is done across scales ranging from *j*_1_ (high frequencies) to *j*_2_ (low frequencies). Thus, the range from *j*_1_ to *j*_2_ should cover the spectral domain of interest with respect to the rhythms to be further extracted. This estimate must also be reliable with respect to the second order statistics involved in (7). There is a compromise between the number of scales to be used and the duration of the epochs in which the statistics are computed. The lowest scales (high frequencies) should not include dominant noisy activity, whereas the largest scales (low frequencies) are limited by epoch duration and edge effects. Our GitHub illustrates the choice of this interval in the algorithm we propose (see *RnB* algorithm).

In summary, our approach bridges the gap between fully time-resolved continuous wavelet transforms (unsuitable for signal processing) and Fourier analysis (unsuitable for time-resolved analysis). The balance between the accuracy of the scale-free model and temporal precision is the key strength of the multi-scale representation of the signal that we introduce.

### Wavelet denoising and extraction of the rhythmic timeseries

To extract the rhythmic activity, wavelet coefficients should be adjusted for the scale-free activity previously described. To do so, we consider the ‘fractional spline wavelet’ bases (Blu and Unser, 2000) to handle the regularity of the original signal, quantified with the parameter *α* = *α*_0_ + *β*^∗^⁄2’ in which *α*_0_ represents the regularity of the rhythmic signal to be extracted and *β* is the scaling exponent of the signal estimated in the previous section. We exploited the property of fractional differentiation of this wavelet family,

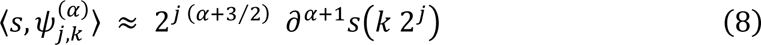

where the partial differentiation operator *∂^r^* corresponds to the product with *f^r^* in the Fourier domain. The following equality can be further shown,

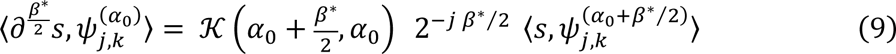

Where

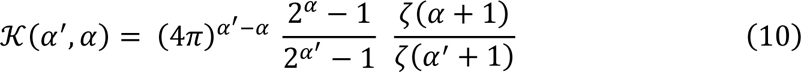

Equation (9) shows that a whitening correction combined with a change of wavelet basis allows to synthetize a time-resolved fractional differentiation, able to reduce the arrhythmicity from the original signal,

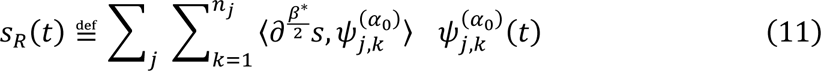

Notice that the fractional differentiation *∂*^*β*^∗/2 of the original signal is precisely the whitening filtering of the power expressed by *Q*(*f*)^−1^ Γ(*f*) previously described in the spectral domain (4). Formula (11) thus expresses this ‘spectral operation’ in the time domain. To summarize, the ‘rhythmic signal’ is defined by

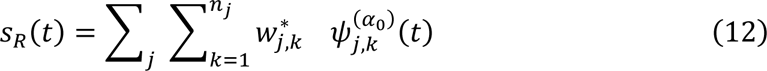

with

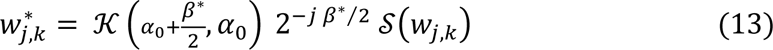

where the *w*_*j*,*k*_’s were computed from the wavelet analysis of the original signal,

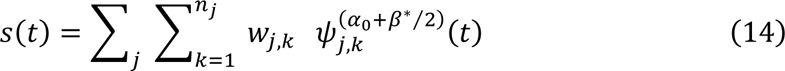

A denoising shrinkage 𝒮 has also been introduced to keep the significant largest wavelet coefficients. It is worth mentioning that each epoch will be analyzed with its appropriate wavelet basis parametrized with a specific *β*^*^ whereas the synthesis is performed in a unique wavelet basis parametrized with *α*_0_. In practice, the regularity of the rhythmic signal is accomplished with *α*_0_ > 2.

The wavelet analysis of an original signal and the synthesis of a rhythmic signal is illustrated respectively on Fig. 2C and Fig. 2E. The discrete wavelet coefficients displayed in Fig. 2C illustrate the stationary gaussian process that governs the wavelet coefficients at each scale. In neural time series, these values are highly influenced by the scale-free process which contributes power in all frequencies. The wavelet coefficients synthetized through fractional differentiation of the original signal are thus corrected for this scale-free process (Fig. 2E).

Although the arrhythmic characteristic of the original discrete wavelet coefficients is attenuated in the discrete wavelet representation of the rhythmic time series (Fig. 2E), the wavelet power of the rhythmic signal does not exhibit the details of the oscillatory content since its representation is driven by scales (Fig. 2F). However, the Fourier power spectrum of the synthetized rhythmic signals will be appropriate to extract the oscillatory components (Fig. 3). The average of the spectral power densities from equivalent rhythmic signal provides an inventory of rhythms that we call ‘spectroscopy’, as illustrated on Fig.3C for epochs collected in sleep. In addition to this spectral signature, we have the distribution of the *β*-exponent that characterizes the arrhythmicity of the rhythm’s background (Fig. 3B).

**Fig.3.**
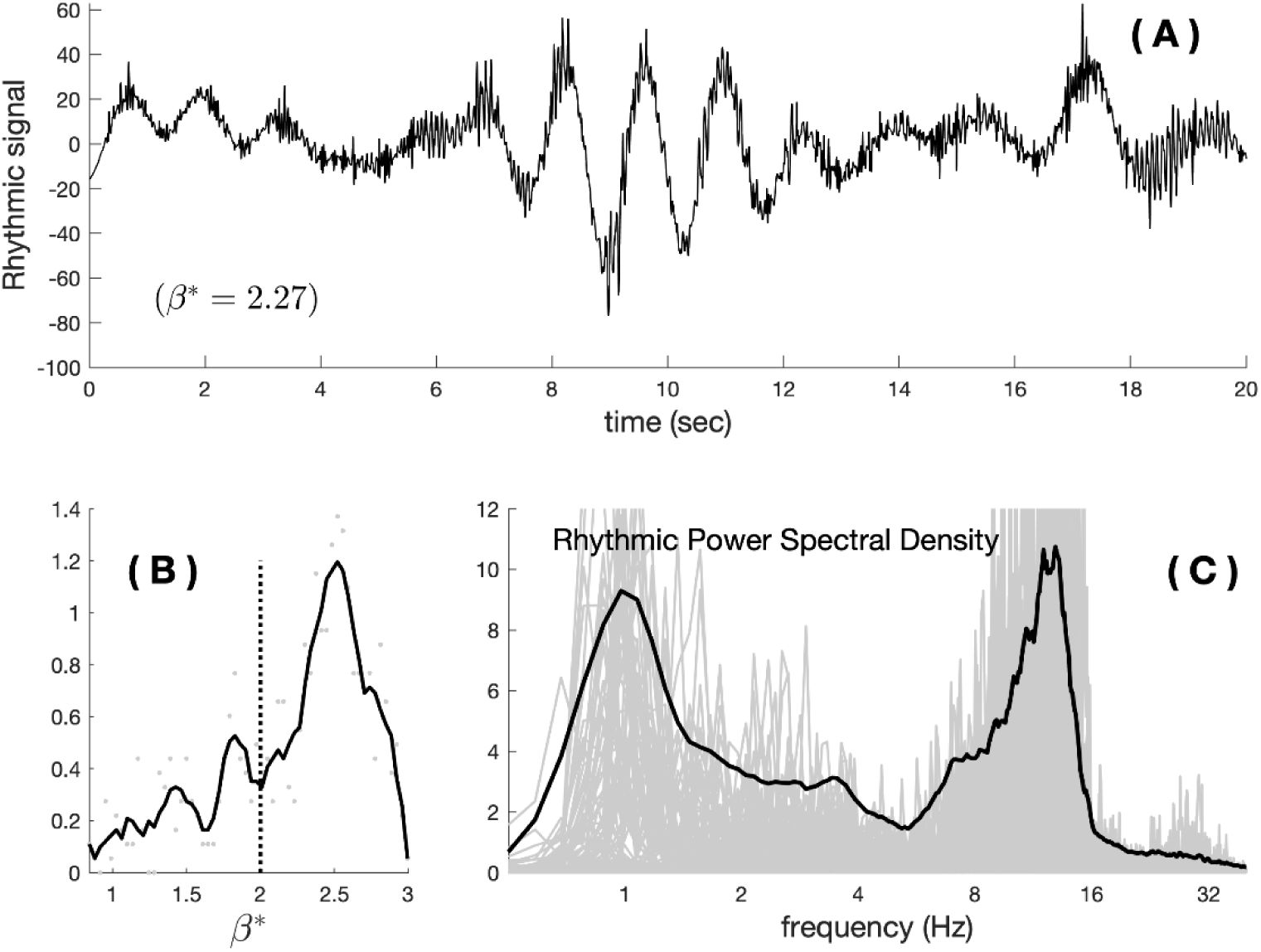
The ‘rhythmic spectroscopy’. (A) The rhythmic signal corresponding to (B) in Fig.2. The value of the *β*-exponent of the removed arrhythmic component is indicated in parenthesis. (B) From 300 epochs in NREM3 from the Anterior Cingulate (see upcoming NREM sleep section), we display the probability distribution of the *β*-exponents. (C) The individual rhythmic spectrums resulting from the Fourier analyses of the rhythmic time series and the ‘rhythmic spectroscopy’ provided with the mean (bold line; see the sleep spectroscopy section for a definition).

### The *RnB* algorithm

In the next sections, the *RnB* algorithm will be used to extract rhythmic timeseries from simulations and intracranial sleep recordings. Description on how to use this algorithm on user-provided data can be found in a publicly available GitHub (https://github.com/michaelcfoti/RnB-Wavelet/). Code for this algorithm is available as a Matlab package licensed by an Apache 2.0 license. The module supports Matlab2021a or later, with dependencies upon the ‘Signal Processing’ toolbox. The package is openly developed and maintained. The project’s repository includes the codebase, a demo which allows to illustrate use of the algorithm on publicly available and user-provided data and the documentation materials. This pipeline allows to process user-provided timeseries, to extract a rhythmic timeserie and perform basic spectroscopy. The rhythmic time series is extracted as a matrix that can be used by the user to perform time-resolved analyses in other pipelines.

Two parameters need to be set to use the algorithm on user-provided data (see the *wRnBMain* script in Github). The parameters linked to the *β*’ estimation relate to the *j*1 and *j*2 scale range to estimate the scale-free process (described in Eq.7).

The parameter *j* refers to the highest scales (lower frequency) upon which the wavelet analysis/synthesis operations are performed to obtain the rhythmic signal, similarly to a high-pass filter. These parameters can be changed according to the input data properties and the documentation on GitHub allows us to understand the impact of such changes. As default parameters used in this work, the scale run from 1 to 9 to determine the arrhythmic *β*′s and the parameter *j* was set to 8.

### Simulation of arrhythmic background and rhythms

Simulations were conducted to assess RnB performance in characterizing background and rhythms. These simulations combined various scale-free backgrounds with transient oscillations. Considering a fractional Brownian motion (fBm) kind of signals, the backgrounds were obtained by filtering a white-noise time series in the frequency domain to obtain a power law 1⁄*f*^*β*^ of the spectral power, while keeping random the Fourier phase. To introduce variability in the background, each simulation trial used a *β*-exponent randomly sampled around a predefined mean value, with a standard deviation of 0.1.

Alpha oscillations were added to these backgrounds using a classical phenomenological neural mass model (Jansen and Rit, 1995; Pinotsis et al., 2012). This model simulates the dynamics of coupled inhibitory and excitatory neuronal populations interacting with a pyramidal cell. The canonical temporal and spectral responses of the model are presented in Fig. 4. These neural mass responses were superimposed onto pure scale-free backgrounds to construct realistic time series.

**Fig.4.**
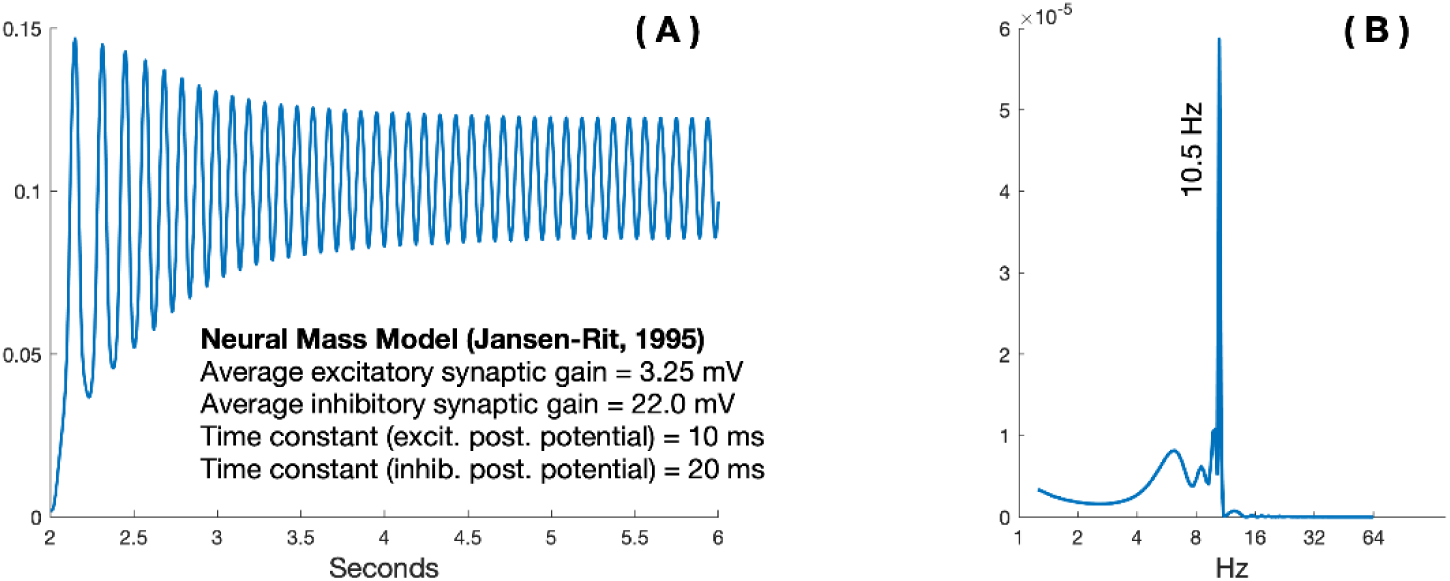
Neural mass oscillation. (A) A four second oscillation produced by a standard neural mass model (with the original parameters). (B) The Fourier power spectrum of the response (A). The alpha peak is accompanied with a lower frequency content that corresponds to the first seconds of the oscillation that does not reach a ‘pure’ harmonic oscillation.

Two batches of simulations were conducted to assess RnB’s accuracy in estimating the maximal rhythmic amplitude from the rhythmic time series and of the scaling exponent.

1. The first batch tested the robustness of RnB estimation for an alpha oscillation (10.5 Hz) with a four second duration simulated at 15 amplitude levels within three arrhythmic background conditions (*β* = 1.7, 2.1 and 2.5).
2. A second batch of simulations assessed the ability of RnB to perform accurate estimations in the context of two concurrent oscillations (delta at 3 Hz delta and alpha at 13 Hz). The amplitude of the delta oscillation was varied across 15 levels and the alpha amplitude remained constant.

For both batches, 300 pure ‘scale-free’ epochs of eight seconds were generated for each amplitude level of the simulated oscillation, with scaling exponent randomly defined around the preceding defined scaling exponents, i.e. 1.7 with a standard deviation of 0.1. Then, 200 randomly chosen time series were selected to incorporate oscillatory activity. Finally, 60 epochs were randomly selected among the 300 epochs per amplitude level for statistical analyses. For each epoch, RnB estimates of maximal rhythmic amplitudes and of the scaling exponents were extracted and included in statistical analyses.

### Statistical testing of the simulation’s parameters

Linear regressions were used to assess the relationship between simulation parameters and *RnB* estimates (estimation of maximal rhythmic amplitude and arrhythmic scaling exponents). Simulations parameters and their interactions were entered as predictors of RnB outcomes in SPSS Statistics 26 software.

### NREM sleep analyses

Our main objectives for NREM analyses were to show that rhythmic time series exhibit characteristics of fundamental NREM sleep rhythms, notably their spectral signature and inter-frequency coupling between delta and sigma rhythms. To do so, we first evaluated how the rhythmic time series extracted from intracranial recordings of NREM sleep provided a reliable representation of cardinal rhythms. Second, we detected sleep slow waves in the rhythmic time series and characterized how these events locally group and synchronize other rhythms in the theta and sigma frequency bands. Finally, we compared delta-sigma coupling between rhythmic and original time series.

### Intracranial recordings

The intracranial recordings used in this work are extracted from the MNI-open iEEG atlas (https://mni-open-ieegatlas.research.mcgill.ca) (Ellenrieder et al., 2020). From this multicentric database providing physiological activity in non-lesional tissues of the sleeping brain, we used the recordings of 16 patients with epilepsy. Analyses were performed on NREM2 and NREM3 sleep recordings and included all the electrodes (200 Hz) localized in the precuneus and the anterior cingulate. All recordings were segmented in 20 seconds epochs. Duration of recordings in each region and sleep stage are described in Table 1.

**Table 1.** Dataset for the intracranial sleep recordings. Total number of 20s epochs and total duration of each sleep stage are shown.

| Region | Sleep stage | # epochs | Total duration (min) |
| --- | --- | --- | --- |
| Precuneus | NREM2 | 967 | 322.33 |
|  | NREM3 | 951 | 317 |
| Ant. Cingulate | NREM2 | 821 | 273.66 |
|  | NREM3 | 809 | 269.66 |

### Spectroscopy of NREM recordings

All NREM sleep epochs were processed with the RnB algorithm to extract rhythmic time series and estimate the distribution of scaling exponents in the time-scale domain. From RnB outputs, we estimated the distribution of scaling exponents and the power spectral density from the rhythmic timeseries separately in NREM2 and NREM3 sleep, what will be referred to as ‘NREM sleep spectroscopy’ (alike Fig. 3B and Fig. 3C). We compared this spectroscopy in each sleep stage to a standard spectral parameterization method (FOOOF). FOOOF was applied on the original intracranial recordings from same NREM epochs that were processed using RnB. For all analyses, FOOOF extracted 3 peaks in the 1-45 Hz range using a fixed aperiodic mode.

### Slow waves detection in the rhythmic signals

We used a proof-of-concept approach to detect sleep slow waves (SSWs) in the rhythmic time series from NREM3 sleep, aiming to demonstrate that the rhythmic time series produced by RnB exhibit the characteristic oscillatory features of SSWs in the time domain.

Rhythmic SSWs (rSSW) were detected in two steps. First, we used a standard SSWs intracranial detection on the rhythmic time series, with an initial delta low-pass filter (0.5-4 Hz), followed by detection of high frequency activity (>32Hz) to define the polarity of the slow waves (Ellenrieder et al., 2016). Second, we performed another detection pass using similar polarity criteria, but now with a band-pass filter adapted to the specific delta found in the spectroscopy of each region and sleep stage (Fig. 7). This adaptive filtering enabled us to focus on the oscillatory content specifically expressed in each region, like recent approaches (Donoghue et al., 2022). The rSSW inventory was defined as the non-redundant union of events detected in both passes. For analyses involving the rhythmic time series (spectroscopy, time-frequency analyses, and PAC), four-second time windows were extracted around the maximal hyperpolarization peak of each rSSW.

For comparative PAC analyses using the original NREM recordings, an additional inventory of SSWs was produced from the original NREM sleep recordings using standard intracranial detection criteria (Ellenrieder et al., 2016) We then extracted four-seconds time windows centered around the hyperpolarisation peak of each detected original SSW.

As in our previous work (Bouchard et al., 2021), we characterized the transition frequency of each SSWs and rSSWs by measuring the time delay (*τ*) between the down phase (hyperpolarization) and up phase (depolarization) maxima. We used an unsupervised Gaussian mixture model (EM algorithm) to estimate each event’s latent ‘switcher’ class (fast or slow) based on the distribution of transition frequencies in each region. Events in the rhythmic signal are referred to as rhythmic slow switchers (rSlowS) and rhythmic fast switchers (rFastS).

### Spectroscopy analyses of rSSWs

We analyzed the spectroscopy of rSSWs during NREM3 sleep. The spectroscopy of all rSSWs was compared to an equivalent number of four seconds NREM3 time windows without detected rSSWs (baseline). Second, we directly compared the spectroscopy of rSlowS to rFastS, as SSWs with a slower transition frequency are known to synchronize NREM sigma rhythms (Nicolas et al., 2022).

### Statistical analyses – impact of the transition frequency on rSSWs spectroscopy

The impact of switcher type on arrhythmic and rhythmic estimates from RnB was assessed using tests computed in SPSS 26. Wilcoxon tests were used to test switcher difference (rSlowS vs rFastS) in the distribution of scaling exponents from RnB. Bilateral t-test were used to test for difference between rSlowS and rFastS for the integrated rhythmic power in frequency bands locally modulated per brain region (delta: 0.5-4 Hz in both regions, sigma: 8-14 Hz in anterior cingulate, 11.5-14.5 Hz in precuneus, theta: 5-9 Hz in precuneus).

### Time-domain analyses of rSSWs

To examine the organization of NREM rhythmic activity around rSSWs, we performed time-frequency analyses (scalograms) around two distinct characteristic markers of rSSWs. For analyses that included all rSSWs (regardless of switcher type), the peak of hyperpolarization was used as the central landmark, consistent with previous studies (Cox et al., 2014). When comparing rSlowS and rFastS specifically, we centered the scalogram on the zero-crossing point between hyperpolarization and depolarization, which represents the midpoint of the transition phase for both types.

We computed two types of scalograms: the standard ‘induced power’ scalogram, which represents the cumulative amplitude of wavelet coefficients and was applied to all rSSWs, and a ‘phase-locked’ scalogram to specifically assess in-phase synchrony of rhythmic signals for switcher comparisons (Tallon-Baudry et al., 1999). For all analyses, we calculated the time-frequency wavelet transform within a one-second window around each central landmark (indicated as t=0 in Fig. 8 and Fig. 9). The scalograms were calculated using the continuous wavelet transform with the Morse analytic wavelet (default parameters: *β* = 20 and *γ* = 3). For visualization only, each scalogram is renormalized using an analytic variant of the Taeger-Kaiser power normalization and each display is individually rescaled between 0 and 1.

### Phase-Amplitude Coupling analyses of rSSWs

To demonstrate that the rhythmic timeseries is sensitive to the phase-coupling of delta and sigma rhythms during NREM sleep, we computed phase-amplitude coupling (PAC) from the original NREM recordings using the SSW inventory and compared it to estimates computed on the rhythmic time series using the rSSWs inventory (rhythmic PAC; rPAC).

For SSWs, we used the phase *φ*(*t*) of the SSW (from the Hilbert transform in the delta band) and the amplitude of the simultaneous sigma activity (10 - 16 Hz) in the original signal. For rPAC, we considered the phase *φ*(*t*) of the rSSW (from the Hilbert transform in the delta band) and the amplitude of the simultaneous sigma activity in the rhythmic timeseries. For rPAC, we adapted the sigma frequency band as defined by the *RnB* spectroscopy in each region (Fig. 7; Anterior Cingulate: 8-14 Hz; Precuneus: 11.5-14.5 Hz).

For each SSW or rSSWs, the PAC is defined by:

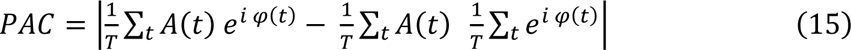

where *T* is the SSW or rSSW duration across which the summation is running.

### Detection of theta bursts associated to rSSWs in the rhythmic timeseries

PAC from the rhythmic timeseries was investigated according to the presence of ‘theta burst’ (TBs) in the precuneus because preceding analyses highlighted that the transition of rSSWs in this region were transiently associated with theta. To do so, we adapted an existing NREM TB detection algorithm to be applied on the rhythmic timeseries (Jiang et al., 2019). We adapted the theta-band filter using our sleep spectroscopy i.e. the theta mode obtained from the rhythmic spectrum in the precuneus (7.02 ± 2.4 Hz in NREM3). The detected events needed to have peaks above ±3 standard deviation of the average power across the epochs with start and stop times defined with ±1 standard deviation. The final TBs inventory kept events with a duration between 0.4 and 1 second. We then extracted 4-sec windows centered around rSSWs co-occurring with TBs (<500ms; rSSWs+TBs events) and around rSSWs that did not (rSSWs-TBs events).

### Statistical analyses of PAC associated with slow waves

PAC were averaged for SSWs detected in the original and rhythmic signal separately for rSlowS and rFastS in each region (anterior cingulate and precuneus). PAC was also investigated according to TB specifically in the precuneus irrespective of switcher type (rSSWs+TBs and rSSWs-TBs).

Mann-Whitney U-tests were used to test for difference in PAC between the original and rhythmic timeseries separately for both type of switchers in the anterior cingulate and precuneus. The impact of switchers type on PAC was also investigated separately in each region for original and rhythmic timeseries separately. Finally, PAC differences between rSSWs according to the presence of a TB (rSSWs+TBs vs rSSWs-TBs) were investigated in the precuneus.

## Results

### Simulations and validation of RnB

We first used realistic simulations to assess *RnB* estimations of both rhythmic and arrhythmic parameters. First, we considered an alpha (10.5 Hz) oscillation in an arrhythmic background (Fig. 5). The signal from this simulation was processed by *RnB* to produce a rhythmic time series (Fig. 5C). The arrhythmic drift which dominated the original time series (Fig. 5A) is removed in the reconstructed signal (Fig. 5C). The spectral density of the rhythmic component provided by *RnB* (Fig. 5D) exhibits a clear signature of the oscillation in comparison with the original Fourier spectrum (Fig. 5B).

**Fig.5.**
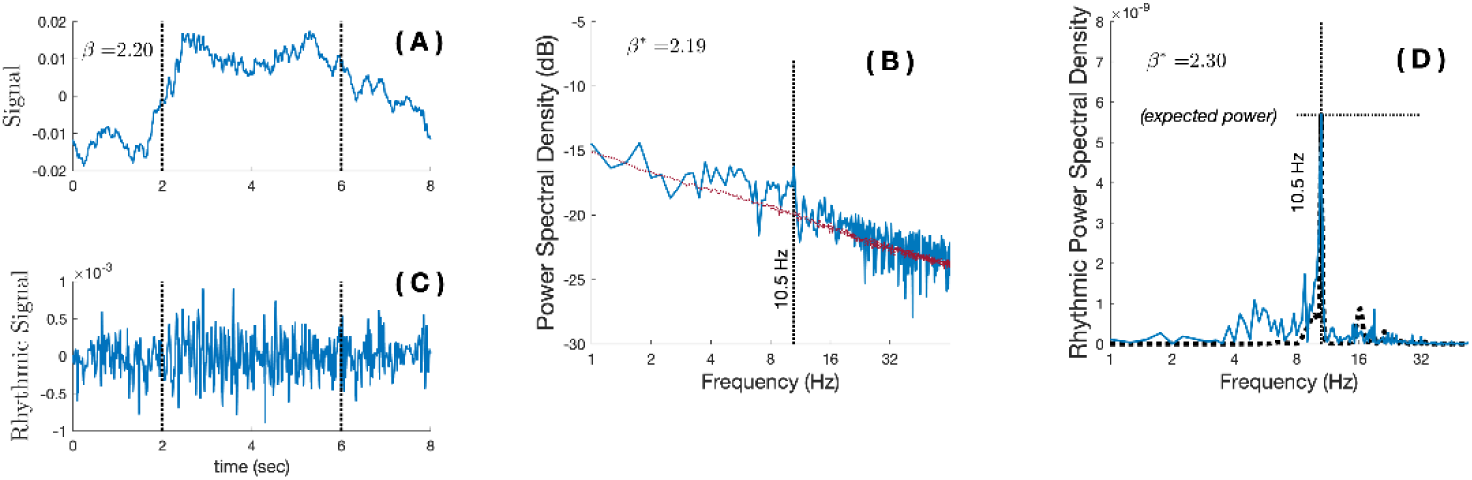
RnB estimation in the wavelet domain. **A.** Example of a simulated time series containing a mixture of a scale-free activity (*β* = 2.2; arrhythmic background) and an alpha oscillation (four seconds 10.5 Hz rhythm). **B.** The Fourier spectra of the original time series *s*(*t*) displayed in (A), mostly dominated by the dominant arrhythmic background. **C.** The output *s*_*R*_(*t*) of the *RnB* algorithm, revealing the rhythmic features of the time series synthesized from the wavelet’s coefficients filtered for the arrhythmic factor. **D.** The Fourier spectra of a single *s*_*R*_(*t*) epoch (in (C)) highlight a clear signature of the simulated oscillation that was added to the arrhythmic background, with a marginal “arrhythmic” contribution. This averaged across multiple epochs define the rhythm’s spectroscopy for further analyses (methods). The black dotted line is the original spectra of the simulated oscillation, without the addition of an arrhythmic background. The ‘expected power’ indicates the maximal power of the simulated oscillation.

Regression analyses were performed to investigate the validity of RnB estimations at different amplitude levels of the simulated oscillations and different arrhythmic background (Fig. 6B). Analyses showed that the rhythmic estimations of the alpha oscillation (amplitude out) scaled remarkably linearly with the amplitude of the simulation (amplitude in) (*b*=0.99, p<0.0001, R^2^=0.98). Rhythmic estimations differed slightly between the three arrhythmic conditions (interaction: t=8.58, p<0.0001; R^2^<0.01), although they remained strongly associated with the amplitude of the simulated rhythm in each of the background conditions (*b*=0.99 to *b*=0.94 from lower to higher scale-free exponents, p<0.0001 in the three arrhythmic conditions).

**Fig.6.**
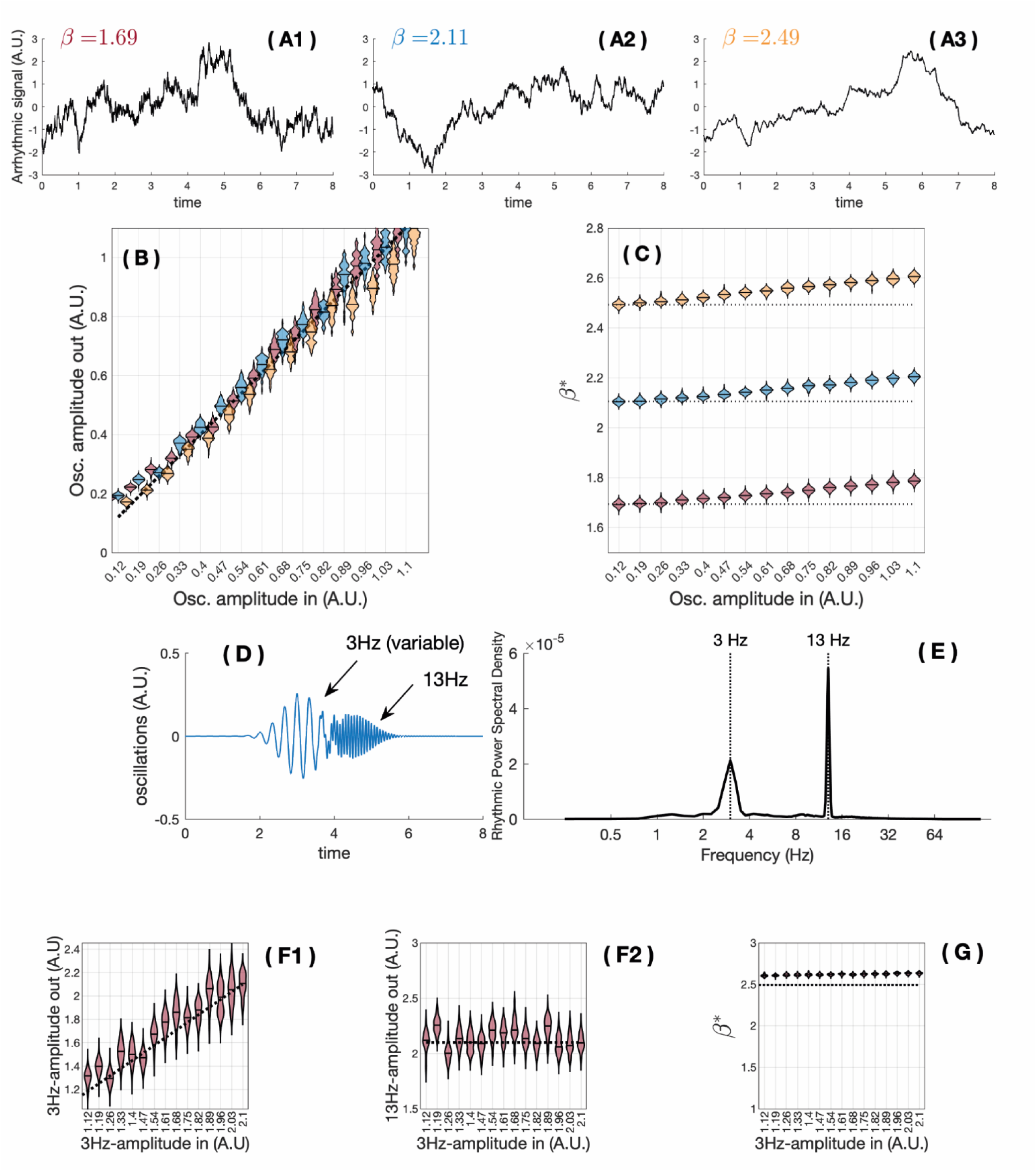
RnB estimation of rhythms and arrhythmic background. (A1, A2, A3). Examples of simulated arrhythmic time series with three different values of arrhythmic background (β) (1.69, 2.11, 2.49); 300 simulations were generated for each arrhythmic condition. For a random set of 200 arrhythmic simulations, an alpha oscillation (10.5 Hz) was added, varying its amplitude at 15 fixed amplitude levels (x-axis on the B plot). For each amplitude level, RnB is applied 60 times on a random selection among all epochs, enforcing a realistic approach where an oscillation may or may not be systematically present. (B). Estimated maximal rhythmic amplitude measured by RnB (Y-axis) plotted against simulated alpha oscillation amplitudes (X axis). Colors of the dots represent the arrhythmic conditions (red: β=1.69; blue: β=2.11; yellow: β=2.49). (C). Estimated scaling exponents out (Y axis) with respect to the simulated alpha oscillation amplitudes (X axis) for the three simulated arrhythmic backgrounds. The dashed line indicates the value in the simulation. (D). An example of rhythmic oscillations to be added to (A3). (E). Spectral signature of the rhythmic component from (D). (F1). Maximal rhythmic amplitude of RnB in the delta range with respect to varying amplitude of this oscillatory component. (F2). Maximal rhythmic amplitude at 13 Hz, varying amplitude of the 3 Hz-oscillation. (G). Response of the RnB algorithm for the scale-free exponent estimation, varying amplitude of the 3Hz-oscillatory component.

Second, we evaluated the accuracy of *RnB* estimates of the arrhythmic background (scaling exponent) and whether they were impacted by the amplitude of the simulated alpha oscillation (Fig. 6 C). As expected, the three simulated arrhythmic background conditions were strongly associated with *RnB* estimates of the arrhythmic background (*β*; change in constant, t=663.3, p<0.0001, R^2^=0.99).

In addition, higher amplitude of the simulated alpha oscillation was associated with higher estimates of the arrhythmic background (*β*; change in slope; b=0.10, t=91.1, p<0.001, R^2^<0.01), although this relationship differed slightly between the three background conditions (interaction; t=9.47, p<0.001; R^2^<0.001; b=0.10 to b=0.11 across the three background conditions).

Third, we investigated whether the presence of a concurrent delta and alpha oscillation affected the accuracy of RnB estimations (Fig. 6 D to G). As expected, the rhythmic amplitude estimation for the delta oscillation (amplitude out) scaled linearly with the amplitude of the simulated delta oscillation (amplitude in; Fig. 6 F1, *b*=0.87, p<0.0001, R^2^=0.85). There was no significant relationship between the amplitude of the simulation delta oscillation (amplitude in) and the estimated maximal alpha amplitude (Fig. 6 F2; *b=*-0.01, p=0.27, R^2^ < 0.01). RnB slightly overestimated the scaling exponent in presence of multiple oscillations, but this does not scale with the amplitude of the simulated delta oscillation (Fig. 6 G, constant=2.7, p<0.001, *b*=-0.03, p=0.31, R^2^=0.001).

Overall, the *RnB* outputs of rhythmic amplitude and scaling exponents are reliable to assess single or concurrent oscillations (alpha and delta), with minor caveats.

First, rhythmic amplitude is slightly underestimated in arrhythmic conditions with high scaling exponents. Second, *RnB* marginally overestimate the scaling exponents when a high amplitude oscillation is present. In both cases, the error from *RnB* is marginal and represents less than 10% of the variance for either of their rhythmic or arrhythmic outputs.

### The spectroscopy of sleep recordings

Given that rhythmic time series provided by RnB are corrected for arrhythmicity, we now demonstrate their potential to characterize genuine NREM sleep rhythms. Our analyses focused on the anterior cingulate and the precuneus, two key regions involved in NREM cardinal delta and sigma rhythms (Murphy et al., 2009; Zerouali et al., 2014).

To better interpret our spectroscopy of NREM sleep recordings, we first illustrated how the NREM sleep power spectrum is decomposed using FOOOF (Donoghue et al., 2020). We then assessed the contribution of the spectroscopy provided by RnB. As expected, NREM2 and NREM3 sleep spectra were characterized by a 1/f arrhythmic background (Fig. 7 A1 and A2). FOOOF confirmed significant sigma peaks in the average NREM spectrum in both stages and regions, with an additional theta peak in the precuneus during NREM2 and NREM3 sleep (Fig. 7 B1 and B2). FOOOF did not reveal peaks below 4Hz, which fits with previous findings indicating the limitations of spectral parametrization methods to identify delta residual spectral power (Gerster et al., 2022).

**Fig. 7.**
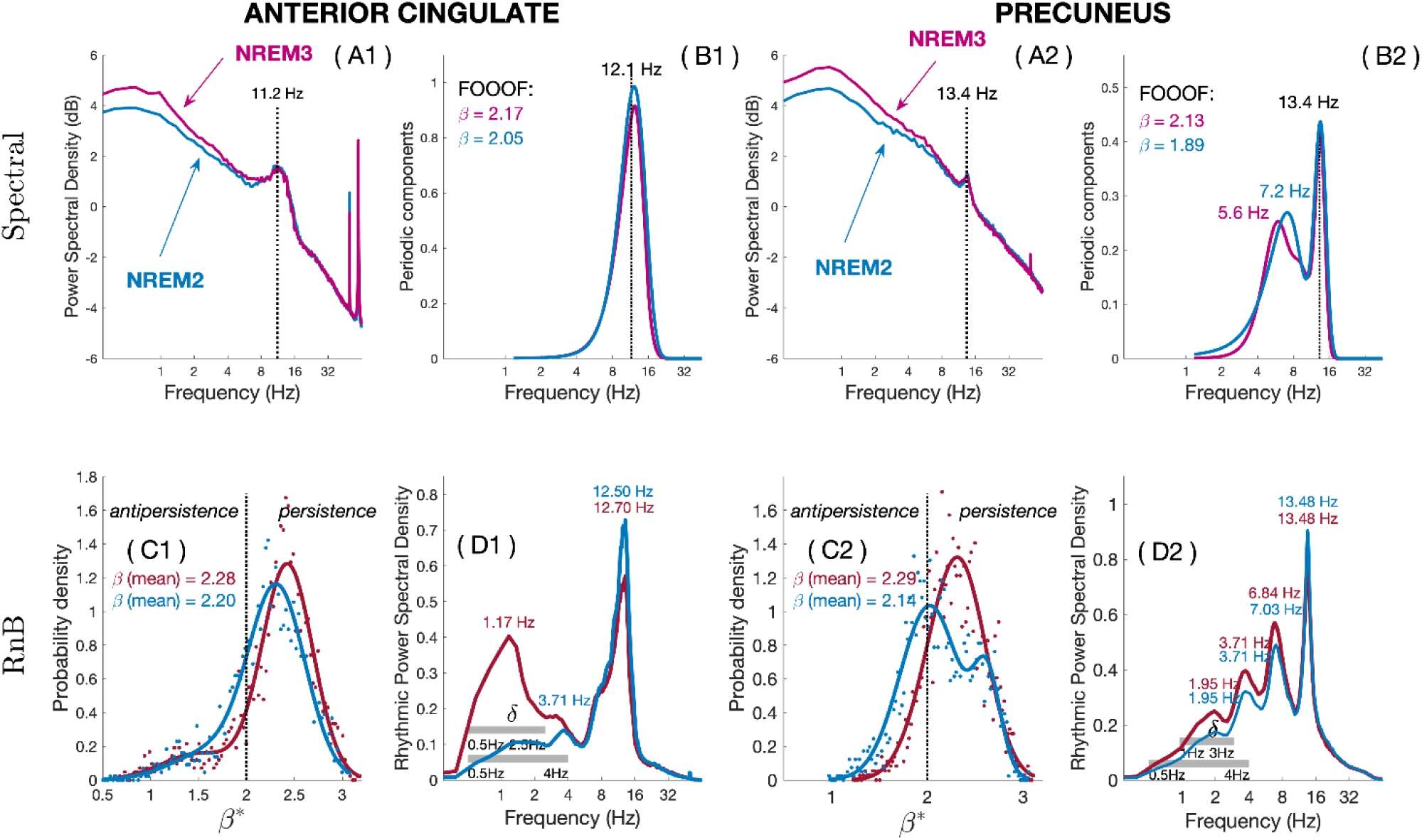
The spectroscopy of NREM sleep recordings. Results from spectral factorization (top row) and the spectroscopy (bottom row) of NREM sleep recordings in the anterior cingulate (two left columns) and precuneus (two right columns). (A1, A2). Standard spectral density per region in NREM2 (blue) and NREM3 (pink) sleep, highlighting a significant arrhythmic 1/f background alongside spectral peaks. (B1, B2). Residual periodic power estimated by FOOOF in NREM2 and NREM3 sleep. Dashed vertical lines indicate found spectral peaks. (C1, C2). Probability density of observed scaling exponents as assessed by *RnB* wavelet coefficients during NREM2 and NREM3. (D1, D2). Spectral analysis from the rhythmic timeseries, highlighting distinctive peaks in each region. Dashed horizontal lines in the delta band indicate the frequency bands to be used for the detection of sleep slow waves in the rhythmic timeseries.

Next, we assessed whether our NREM sleep spectroscopy provides a reliable representation of NREM cardinal rhythms. Interestingly, the NREM sleep spectroscopy highlighted higher delta power in NREM3 as compared to NREM2 sleep, although the specific frequency bands varied between the two regions (Fig. 7 D1 and D2). In the anterior cingulate, a low-frequency delta rhythmic peak (1.17*Hz*) predominated in NREM3 as compared to NREM2 sleep, whereas two delta rhythmic delta peaks (1.95 and 3.71 Hz) were found in the precuneus during

NREM3 and NREM2 sleep. A rhythmic theta peak (≈ 7*Hz*) was also found specifically in the precuneus during NREM2 and NREM3 sleep. Additionally, rhythmic sigma peaks were found in both regions, with slightly faster frequencies in the anterior cingulate (12.5 Hz) as compared to the precuneus (13.5 Hz), and lower maximal power in NREM3 compared to NREM2. *RnB* also provided distributions of scaling exponents *β* in each region (Fig. 7 C1 and C2), with higher scaling exponents (higher persistence) in NREM3 as compared to NREM2 sleep in both regions. Thus, RnB spectroscopy reveals a more exhaustive NREM sleep spectral content as compared to FOOOF.

### Rhythmic sleep slow waves modulate rhythmic and arrhythmic activity

Given that delta activity was higher in NREM3 in both regions, we detected sleep slow waves (SSWs) in the rhythmic time series (rSSWs), cardinal NREM rhythms, and assessed how they modulate other rhythms during NREM3 sleep. The analysis of rSSWs showed an expected biphasic shape, with a suppression of higher frequency during their downstate (Fig. 8 A1 and A2). Interestingly, time-frequency analyses of the rhythmic timeseries associated with rSSWs in NREM3 showed that rSSWs in the anterior cingulate were specifically associated with sigma activity in their depolarization period during NREM3 sleep (Fig. 8 B1 and B2).

**Fig. 8.**
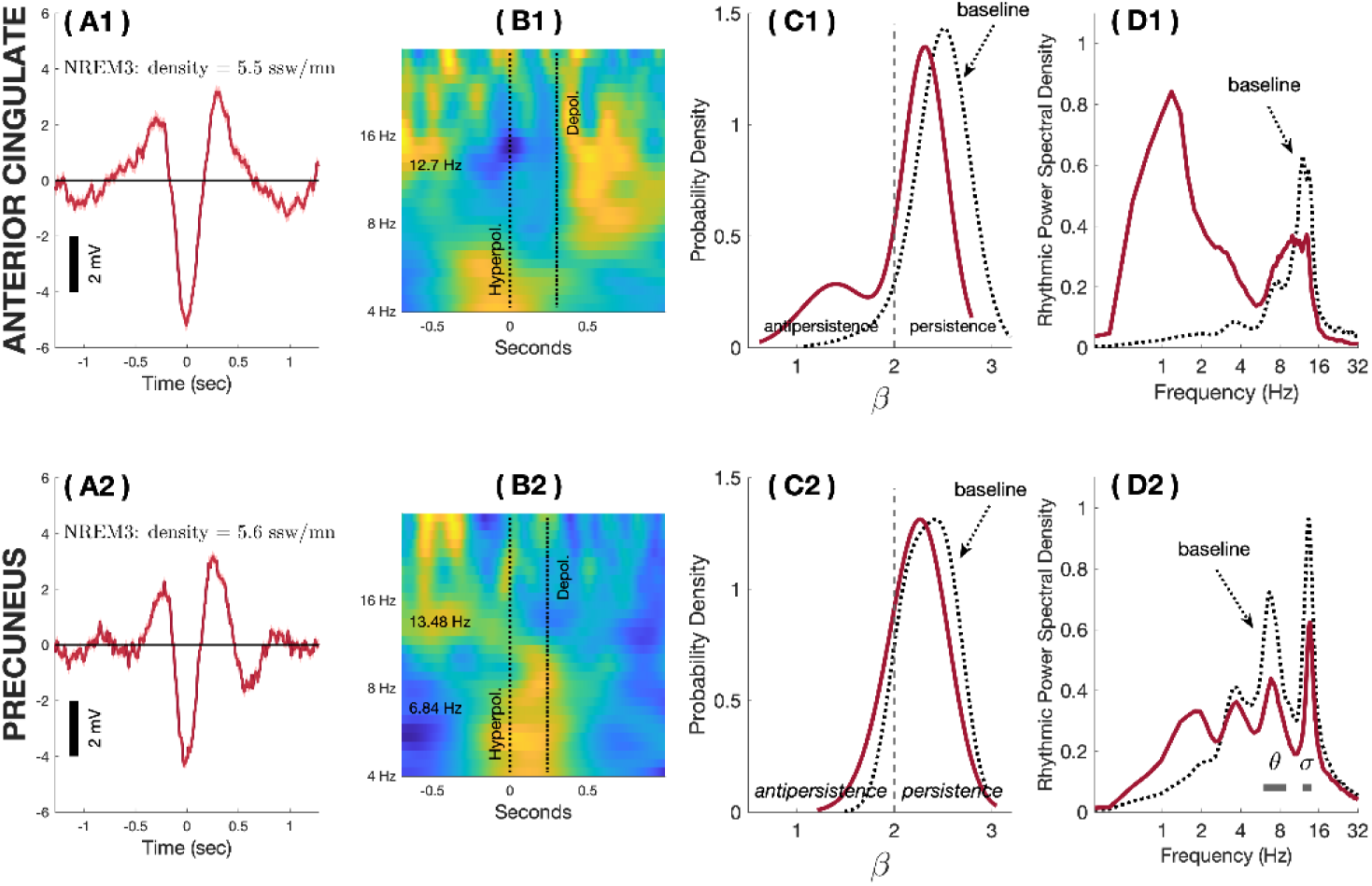
SSWs in rhythmic time series from the anterior cingulate and precuneus during NREM3 sleep. (A1, A2). Average waveforms of rSSWs detected during NREM3 in the anterior cingulate (top) and precuneus (bottom). (B1,B2). Induced scalograms of the rhythmic time series associated with SSWs coregistered with respect to the hyperpolarization phase. Red/yellow indicate higher normalized power than blue. (C1,C2). Distribution of scaling exponents across epochs with rSSW (plain) and without rSSW (dashed lines). (D1,D2). Spectroscopy of epochs with rSSWs (plain) and without rSSWs (dashed lines) during NREM3 sleep.

Spectroscopy-based analyses of rSSWs were also performed to assess how rSSWs transiently modulate rhythmic and arrhythmic NREM activity. As compared to baseline NREM epochs (epochs without rSSWs), most epochs with rSSWs had lower scaling exponents (lower persistence), although this effect was predominant in the anterior cingulate (Fig. 8 C1 and C2). Epochs with rSSWs also showed higher delta and lower sigma rhythmic power than baseline epochs in both regions, although the anterior cingulate showed a stronger increase in rhythmic delta and a wider rhythmic sigma (Fig. 8 D1 and D2). The theta peak previously observed in the precuneus also showed lower power during rSSWs as compared to the baseline.

### The spectroscopy of SSWs differ according to their transition period

SSWs can be categorized as slow or fast switchers, depending on the duration of the *transition period* between their hyperpolarization and depolarization phase (Bouchard et al., 2021). Interestingly, sigma rhythms are preferentially nested in the transition of SSWs with longer transition periods (slow switchers), as compared to fast switchers during NREM sleep (Nicolas et al., 2022). We thus evaluated whether sigma activity after rSSWs was due to slow switchers.

We categorized each rSSW during NREM3 as a rhythmic slow switcher (rSlowS) or rhythmic fast switcher (rFastS; methods; Fig. 9). We reproduced the clustering of rSSWs between rSlowS and rFastS in the rhythmic time series, with a distribution skewed toward rFastS in both regions (Fig. 9 A1 and A2). The spectroscopy of rSlowS differed from rFastS particularly in the anterior cingulate. In this region, rSlowS showed a bimodal distribution of scaling exponents with persistent and antipersistent modes, whereas rFastS showed a distribution of scaling exponents with a clear persistent mode (Fig. 9 C1; Wilcoxon test : z-statistic: -4.49, p<0.001). Compared to rFastS, rSlowS also showed significantly higher rhythmic delta and higher rhythmic sigma power in the anterior cingulate (Fig. 8 D1, t-test p<0.05). In the precuneus, arrhythmic and rhythmic estimates were superposable between rFastS and rSlowS (Fig.9 C2 and D2).

**Fig. 9.**
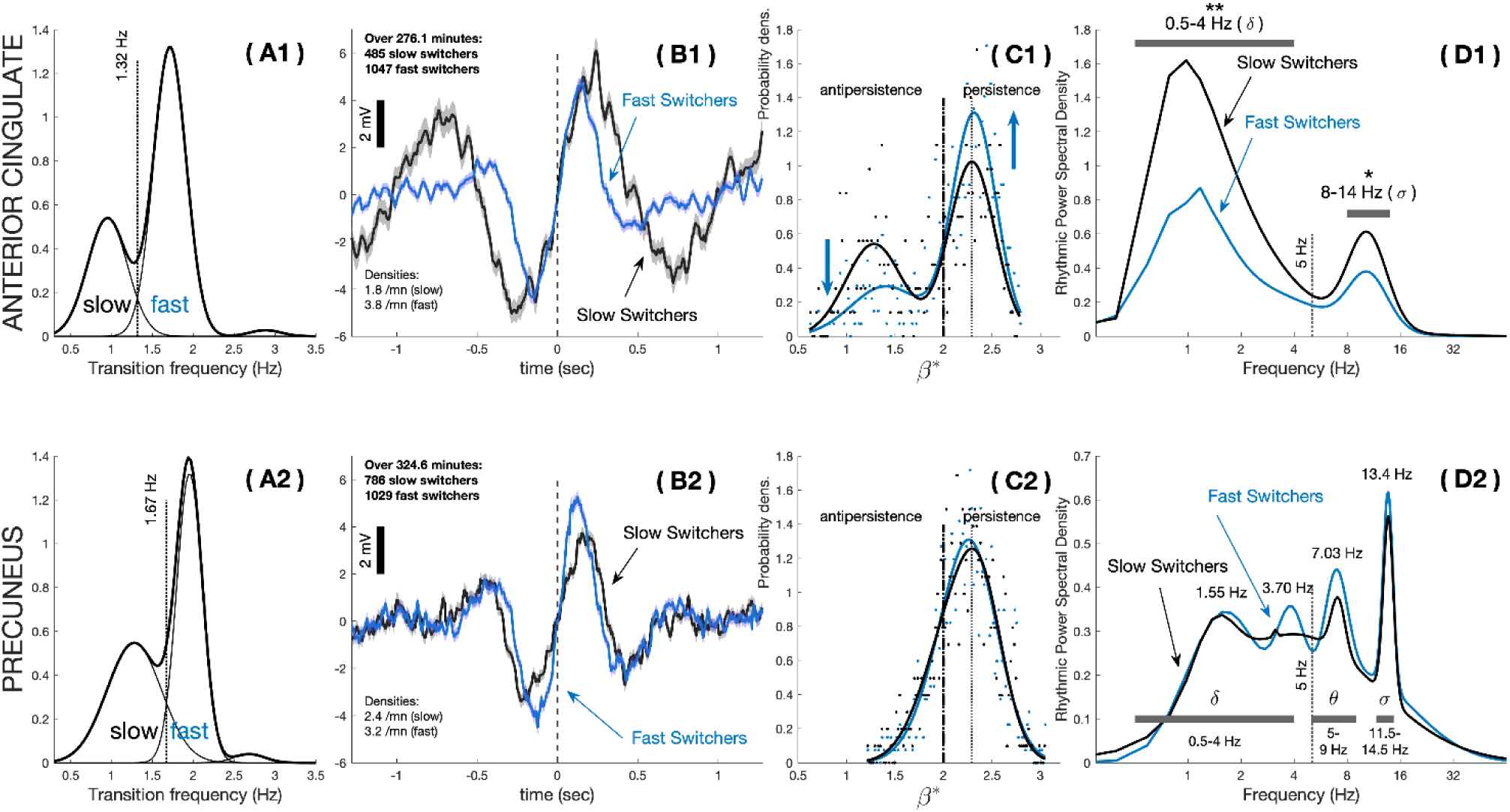
The spectroscopy of slow (rSlowS) and fast switchers (rFastS) during NREM3. (A1, A2). The probability density of rSSWs according to their transition frequency is shown for the anterior cingulate (A1) and precuneus (B1), highlighting a bimodal distribution in both regions. The transition frequency to cluster rFastS and rSlowS is shown as a dashed vertical line. (B1, B2). Average waveform of rSlowS and rFastS, with respect to the separation frequency are defined in (A1) and (A2) respectively. Averages are centered around the zero-crossing markers. (C1, C2). Probability density of the scale-free exponents is estimated by *RnB* for rSlowS and rFastS in each region. (D1, D2). Spectroscopy of the rhythmic switchers in NREM3 are shown for both regions. The horizontal bold vertical bars depict local frequency bands which were considered for further PAC analyses. Bilateral T-tests were performed in each region to investigate switcher-related differences in the integrated rhythmic power for each region (anterior cingulate: delta between 0.5 and 4 Hz, sigma between 8-14; precuneus: delta between 0.5 and 4 Hz, theta between 5-9 Hz, and sigma between 11.5 – 14.5 Hz). Significance is depicted using stars (p-values: * <0.05; **<0.01).

### rSlowS and rFastS regroup differently transient sigma and theta rhythms

To evaluate how rSSWs synchronize NREM rhythms with respect to their transition period, we assessed the scalograms of rSlowS and rFastS derived from the rhythmic time series (Fig. 10). Induced and phase-locked rhythmic power were both considered to assess how rSSWs group or synchronize rhythmic activity, respectively. As sigma rhythms occur specifically during the transition from hyperpolarization to depolarization particularly for slow waves with slower transition (Nicolas et al., 2022), we hypothesized that rSlowS would show higher phase-locked sigma power during their transition as compared to rFastS.

**Fig.10.**
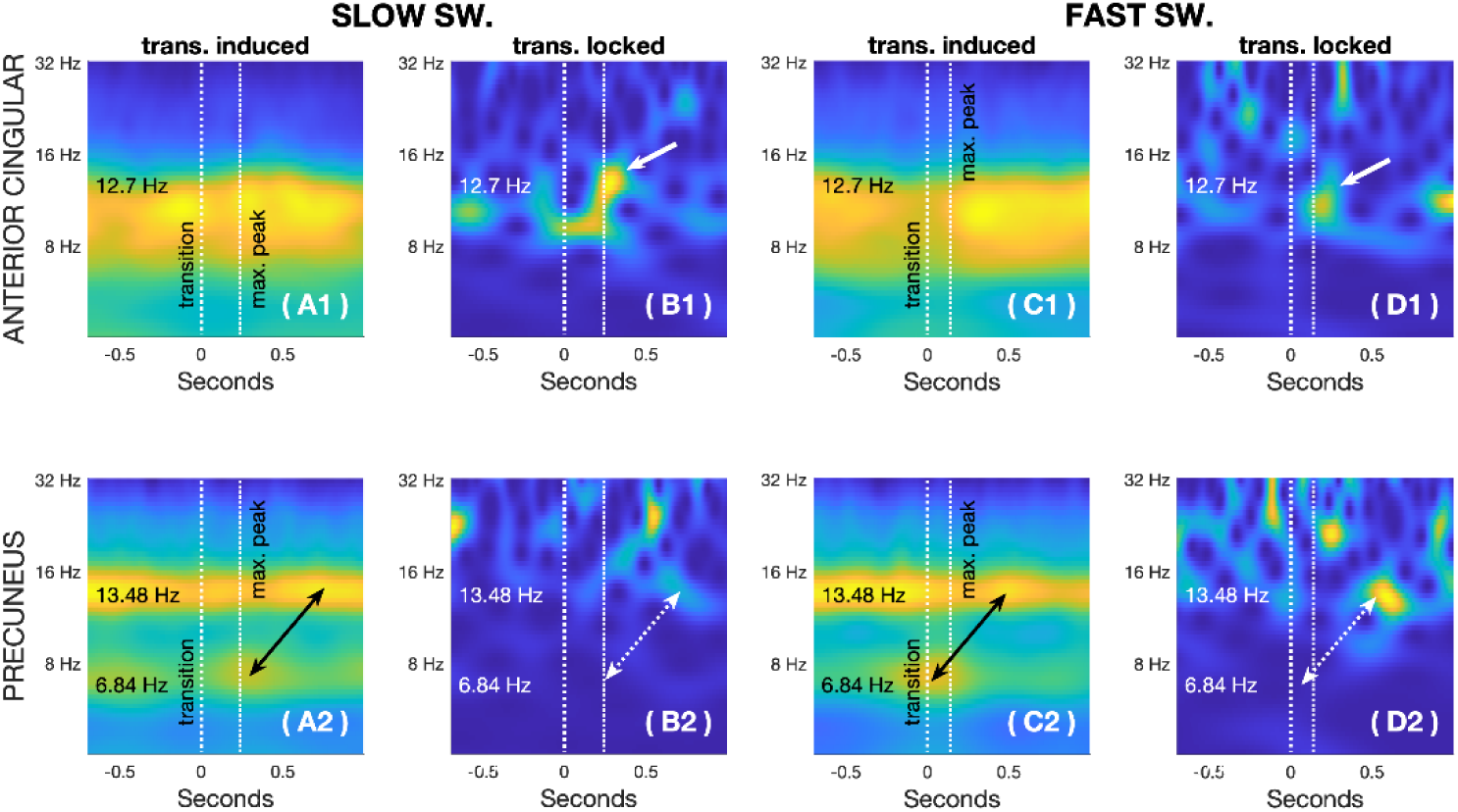
Induced and phase-locked rhythmic power synchronized with the transition of rSlowS and rFastS during NREM3 sleep. Scalograms of induced (A1, A2, C1, and C2) and phase-locked (B1, B2, D1, and D2) power for the rSlowS (Left panels) and the rFastS (Right panels), co-registered around their zero crossing, in the anterior cingulate (top panels) and the precuneus (bottom panels). Vertical lines indicate the transition markers for rSlowS and rFastS (zero-crossing in bold; the depolarization ‘max. peak’ as estimated from the averaged signal in Fig.9 B1 and B2). The arrows in top panels (anterior cingulate) indicate phase-locked power around the transition of rSlowS, whereas arrows in bottom panels (precuneus) emphasize the temporal relationship between the induced theta and phase-locked sigma activity predominantly for rFastS. Each plane is individually normalized for visualization (hot colors for higher power).

Induced sigma power was observed across rSlowS and rFastS in both regions (Fig. 10 A1, A2, C1, C2). The precuneus also showed induced theta power around the transition phase of rFastS and rSlowS (Fig. 10 A2, C2). As expected, rSlowS in the anterior cingulate showed phase-locked sigma rhythmic power in their transition, with increasing frequency and reaching maximal power synchronously with their depolarizing peak (Fig.10 B1). Although induced ‘theta bursts’ (TBs) in the precuneus were not sufficiently synchronized around the transition to produce a theta signature in the locked scalogram, phase-locked rhythmic sigma power occurred later with a delay around 400 msec preferentially for rFastS (Fig.10 B2, D2).

### rSlowS and TBs promote SSW-sigma coupling specifically in rhythmic time series

To evaluate how sigma amplitude is systematically modulated by the phase of delta rhythms in NREM sleep, we computed delta-sigma phase-amplitude coupling (PAC) during SSWs in NREM3. We considered PAC from SSWs in the original EEG signal and from rSSWs detected in the rhythmic time series (rhythmic PAC: rPAC). rPAC was computed using delta frequencies which stood out locally in each region’s spectroscopy (Fig. 7, see methods), whereas PAC was computed using standard SSW (0.5-4Hz) and sigma (12-16Hz) frequencies. We hypothesized higher PAC for rSlowS and rFastS detected in the rhythmic time series as compared to events detected in the original signal since arrhythmic interferences should be minimized.

As shown in Fig. 11 (A and B), rPAC values were significantly higher than PAC values for both types of switchers in the two brain regions (Mann-Whitney U-tests: p < 0.001 for all events). These results suggest that phase interferences are reduced in the rhythmic time series, revealing higher phase coupling between delta and sigma rhythms. Additionally, rPAC was significantly higher for rSlowS as compared to rFastS in the anterior cingulate during NREM3 (Fig. 11 A, U=2.82, p=0.005), as expected. This difference between SlowS and FastS in the anterior cingulate was not significant with standard PAC (U= 1.85, p-value=0.06). In the precuneus, no significant differences in rPAC were found between switchers (Fig. 11 B, U=0.18, p>0.1).

**Fig.11.**
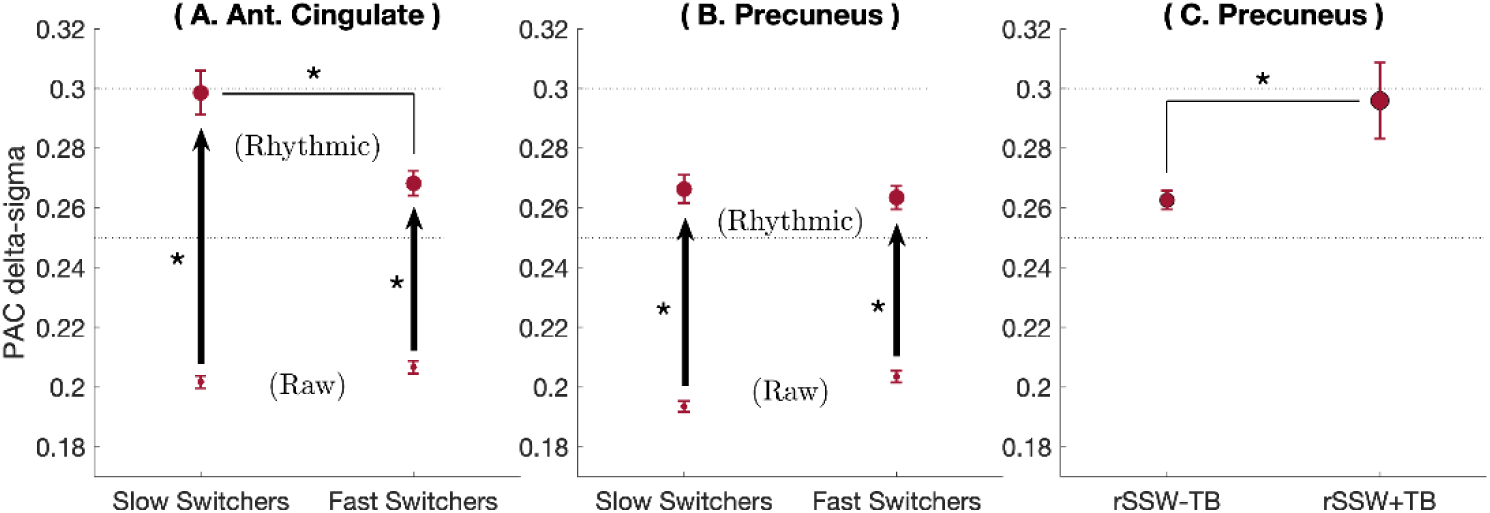
Delta-sigma PAC from original and rhythmic time series. Average values and standard error of the mean for rPAC (rhythmic) and PAC (raw), for slow and fast switchers during NREM3 sleep in the anterior cingulate (A) and in the precuneus (B). In the precuneus, rPAC values are shown in (C) for all rSSWs without TBs (rSSW-TBs) and for rSSWs co-detected with a TBs (rSSW+TBs). For each event type, arrows indicate significant differences between PAC and rPACs values. Stars indicate significance value (*: p≤0.01).

Since previous literature suggests that NREM TBs precede slow-wave-spindle complexes and facilitate delta-sigma PAC (Gonzalez et al., 2018), we explored the effects of TBs on rPAC during rSSWs in the precuneus. This analysis is prompted by the specific theta peak and induced theta found in the precuneus spectroscopy and with the transition of rSlowS and rFastS in the precuneus scalogram analyses (Fig.10). We designed a rhythmic TBs detector that used the theta peak found in the precuneus spectroscopy. Then, we identified rSSWs co-occurring with TBs (<500ms; rSSWs+TBs events) and rSSWs that did not (rSSWs-TBs). We hypothesized higher delta-sigma rPAC for rSSWs+TBs events as compared to rSSWs-TB in the precuneus.

As shown in Fig. 11 C, delta-sigma rPAC was higher for rSSWs+TBs compared to rSSWs-TBs (U=2.46, p=0.01). Importantly, rSSW+TBs events represented around 6% of all rSSWs in NREM3 (116/1815 rSSWs). Although rSSW+TBs are rare, they constitute a significant modulator of rPAC in NREM3. Thus, the rhythmic time series shows globally higher delta-sigma coupling. Furthermore, rPAC was higher for rSlowS in the anterior cingulate and was significantly amplified with TBs in the precuneus.

## Discussion

Interference between background and rhythms in the time domain is a challenge that current spectral parametrization methods cannot address. We introduce a wavelet-based method able to disentangle the rhythmic time series from the arrhythmic background. We showed that rhythmic time series adequately extract rhythmic activity in either simulations or in NREM sleep, with enhanced coupling estimates.

RnB allows time-domain analyses of brain rhythms free from arrhythmicity, whereas spectral methods focus on properties of “periodic” peaks and aperiodic background (Wen and Liu, 2016; Donoghue et al., 2020). Usually characterized as a 1/f slope (i.e., a scaling exponent) varying over time, arrhythmicity indicates fluctuations of scale-free brain dynamics (He et al., 2010; Miskovic et al., 2019). Pure arrhythmic EEG (i.e., without cardinal oscillations) represents around 80% of the recording time in NREM2 and more than 50% in NREM3 sleep (Helfrich et al., 2021). This ubiquitous background also interferes with spectral and temporal estimates of brain rhythms: time series with higher scaling exponents are characterized by slower fluctuations contributing to delta power in standard spectral analyses which may, for instance, be confounded for sleep slow waves (Donoghue et al., 2022; Gerster et al., 2022). As previously reported (Lendner et al., 2020; Bódizs et al., 2021; Horváth et al., 2022), our results showed higher scaling exponents in deeper NREM sleep. We also showed that scaling exponents fluctuate across sleep epochs, which was previously reported in multifractal analyses (Lina et al., 2019). This overnight variability in arrhythmicity may reflect time-dependent changes in multiple neurophysiological parameters during sleep, such as the excitation-inhibition balance (Niethard et al., 2016; Gao et al., 2017) and the dynamics of brain networks (Brake et al., 2024). So far, amplitude and phase estimates of oscillatory activity remained unaccounted for arrhythmicity. *RnB* allows to extract phase and amplitude estimates of rhythmic activity underlying genuine NREM sleep oscillations in the time domain, controlling for ‘instantaneously’ differences in the arrhythmic backgrounds.

Recent simulation studies pinpointed limitations in spectral methods to reveal rhythmic delta due to interferences between arrhythmicity and low frequencies in the Fourier domain (Gerster et al., 2022). Hence, previous sleep studies using such methods mostly focused on sigma periodic peaks (Bódizs et al., 2021; Schneider et al., 2022). Interestingly, our simulations showed that *RnB* robustly estimate concurrent delta and sigma rhythms across different scale-free scenarios. Furthermore, *RnB* detected delta, theta, and sigma differences between NREM2 and NREM3 in the anterior cingulate, which correspond to changes expected in NREM sleep (Steriade, 2006). We argue that RnB allows an accurate spectroscopy of NREM rhythms due to the wavelet based ‘punctual’ estimations and corrections in the time-scale domain. This allows to synthesize rhythmic delta activity in the time-scale domain, which otherwise overlaps with the scale-free process in the Fourier domain.

Compared to the precuneus, the anterior cingulate spectroscopy showed stronger differences in arrhythmic and rhythmic activities between NREM2 and NREM3 sleep. Specifically, rhythmic timeseries from the anterior cingulate showed higher rhythmic delta power in NREM3 as compared to NREM2 sleep, consistent with the well-known anterior predominance of NREM delta power and of sleep slow waves (Vyazovskiy et al., 2007; Hubbard et al., 2020). This facilitated the detection of sleep slow waves (SSWs) in the rhythmic time series (rSSWs) from NREM3 sleep using spectroscopy-adapted criteria, aligning with recent guidelines to adapt detection criteria to local frequency content (Muehlroth and Werkle-Bergner, 2020; Donoghue et al., 2022).

rSSWs in the anterior cingulate were associated with higher rhythmic delta power and lower scaling exponents compared to baseline. Lower scaling exponents during rSSWs in the anterior cingulate resonate with a recent study reporting a flatter slope (lower scaling exponent) during the depolarization phase of SSWs in anterior regions, suggesting higher excitation in this phase (Lendner et al., 2020). This is consistent with the anterior predominance of the slow oscillation, during which neurons are strongly depolarized in the active phase (Steriade, 2006). Furthermore, rSSWs modulated sigma power differentially the anterior cingulate and precuneus, consistent with previous EEG findings on SSWs (Cox et al., 2014). This anterior specificity may reflect the cortico-genesis of SSWs, as these waves originate in the anterior cingulate before spreading in other regions (Murphy et al., 2009). Thus, we hypothesize that rSSWs, expressed above the arrhythmic background, result from the underlying slow oscillation during NREM sleep.

We reproduced the ‘fast-slow’ switchers classification of the rSSWs observed in scalp EEG sleep recordings (Bouchard et al., 2021). In the anterior cingulate, rSSWs with slower transitions (rSlowS) displayed distinct characteristics compared to those with faster transitions (rFastS), including higher delta and sigma rhythmic power. These differences align with evidence that SSWs with longer depolarization periods show larger amplitudes, higher sigma synchronization and sleep spindle occurrence (Botella-Soler et al., 2012; Cox et al., 2018, 2020; Chylinski et al., 2022). Furthermore, rhythmic sigma power was phase-locked specifically after rSlowS transition in the anterior cingulate, and these events exhibited higher delta-sigma rPAC compared to rFastS. Together, these findings suggest that rSlowS from the anterior cingulate play a pivotal role in mediating interactions between rSSWs and NREM sigma rhythms.

Supporting this, previous findings indicated that sleep spindles in anterior regions did not show a clear relationship to fast switchers and preferentially occurred during the depolarization phase of a subset of slow switchers (Chylinski et al., 2022). Consistently, we observed a bimodal distribution of scaling exponents specific to rSlowS in the anterior cingulate, with a subset exhibiting lower values (i.e. antipersistence). Recent work suggests that such arrhythmic properties reflect local network dynamics involved in long-range connectivity (Brake et al., 2024). Interestingly, slow switchers coupled with sleep spindles are known to show higher frontal EEG connectivity (Bouchard et al., 2021). Thus, we hypothesize that the antipersistent arrhythmic regime linked to rSlowS in the anterior cingulate may contribute to the emergence of cortical networks associated with sleep spindles.

Interestingly, while the coupling between rSSWs and sigma in the anterior cingulate depended on the transition frequency of rSSWs, it relied instead on the co-occurrence of TBs and rSSWs in the precuneus. TBs have been identified as an “early” down state when the slow oscillation is less synchronized, facilitating subsequent SSWs up states and the later occurrence of sleep spindle (Gonzalez et al., 2018; Jiang et al., 2019). Thus, TBs could help to drive sigma synchronization in posterior regions, where the cortical slow oscillations are usually less synchronized. Furthermore, we can hypothesize that transient sigma rhythms associated to rTBs or rSlowS could synchronize across brain regions to form “global” spindles. Interestingly, EEG “global” spindles involve progressive synchronization of MEG sigma rhythms from the precuneus to the anterior cingulate (Zerouali et al., 2014). Thus, transient rSlowS and TBs could be local cortical features contributing to sigma synchronization during global spindles.

Delta-sigma PAC was higher for SSWs detected in the rhythmic as compared to the original signal. A few reasons could account for this. First, it is possible that the spectroscopy-improved rSSW detection criteria increased the representation of genuine slow oscillations, which led to better disclose delta-sigma coupling (Aru et al., 2015). The rhythmic time series might also better estimate the inherent phases of the cardinal rhythms, as arrhythmic interferences have been attenuated (Donoghue et al., 2022). In any case, it is likely that coupling in the rhythmic time series provides a better insight into the oscillatory-dependent hippocampo-thalamo-cortical interactions altered in numerous disease (Helfrich et al., 2019; Mendes et al., 2021). Currently, PAC metrics express high heterogeneity and are often low, preventing their clinical use (Mander et al., 2017; Jurkiewicz et al., 2021). By jointly improving the detection of NREM rhythms and the assessment of their coupling, *RnB* could decrease the heterogeneity of PAC or phase-based findings.

*RnB* represents a significant paradigm shift in EEG signal processing. It constitutes a crucial step towards disentangling electrophysiological rhythms from arrhythmicity in brain time series. However, further refinements could help to estimate rhythmic processes even more accurately. Specifically, our current rhythmic signal may not be considered as a “sparse representation” in terms of oscillatory events and future work could improve its classification. For instance, instead of using standard detection criteria into the rhythmic time series, previously proposed matching pursuit methods could be applied to the rhythmic signal to better identify basic patterns corresponding to NREM rhythms (rSSWs, TBs, sigma rhythms). This would allow the development of detectors relaxing the need of prior detection criteria (Durka and Blinowska, 1995; Krstulović and Gribonval, 2006; Durka et al., 2024). Alternatively, previously proposed cycle-by-cycle analysis could also be used directly in the rhythmic signal to identify periods where such events manifest. Thus, the application and refinement of our ‘*sleep spectroscopy’* could be crucial to provide updated detection criteria for NREM sleep transient events across the lifespan and clinical conditions.

